# A modelling technique unifying four paradigms of metacommunity theory

**DOI:** 10.64898/2026.04.21.719942

**Authors:** Sara Shahin, Jacob D. O’Sullivan, Axel G. Rossberg

## Abstract

Metacommunity theory explains how species distributions arise from local population dynamics and dispersal between habitat patches. Four conceptual paradigms—patch dynamics, species sorting, mass effects, and demographic stochasticity—have emerged as frameworks for understanding metacommunity structure and dynamics, but their integration remains an open problem. Here we introduce a probabilistic-stochastic-deterministic (PSD) modelling framework that unifies these paradigms within a single mathematical description. PSD approximates individual-based models (IBM) with computational efficiency comparable to ordinary differential equations (ODE) while capturing demographic stochasticity and permitting analytical treatment. Through validation against IBM simulations in single-patch communities and spatially explicit metacommunities with rock-paper-scissors dynamics, we demonstrate that PSD accurately reproduces IBM behaviour where ODE models fail, specifically when demographic stochasticity dominates during immigration. For metacommunities with long-distance dispersal, we analytically derive the period of a slow collective oscillations, revealing body-mass and dispersal-rate dependencies invisible to ODE theory. Our analysis shows that the four paradigms represent valid descriptions in different regions of parameter space, controlled by individual body-mass, immigration rate, and regional species richness. The PSD framework thus provides both a practical simulation tool and an analytical machinery for predicting metacommunity dynamics across ecological regimes.

## 1 Introduction

Metacommunity theory provides a framework for understanding how species distributions arise from the interplay of local population dynamics and dispersal between spatially discrete habitat patches (Leibold et al. 2004; Holyoak et al. 2005). It bridges the ecology of the local community, where species interactions and environmental filtering operate, and regional biogeography, where dispersal limitation shapes species pools (Ricklefs 2008; Logue et al. 2011; MacNeil et al. 2009).

At the most fundamental level, metacommunities can be described by individual-based models (IBMs) that track how individuals reproduce, interact, die, and disperse between patches (Grimm and Railsback 2005; DeAngelis and Grimm 2014). IBMs capture the full complexity of ecological dynamics, including demographic stochasticity arising from discrete birth and death events. However, IBMs provide limited analytical insight into emergent macroscopic patterns. To address this, researchers have introduced simplifying abstractions, giving rise to four conceptual paradigms (Leibold et al. 2004): *patch dynamics*, emphasising stochastic patch colonisation and extirpation events; *species sorting*, emphasising deterministic responses to environmental heterogeneity and biotic interactions; *mass effects*, emphasising dispersal-maintained sink populations; and *demographic stochasticity*, emphasising chance events at low population sizes. These paradigms have often been treated as competing alternatives (Cottenie 2005; Winegardner et al. 2012; Driscoll and Lindenmayer 2010), yet their common origin in individual-based dynamics suggests that a unifying synthesis should exist that recovers each paradigm as a limiting case within a more general description (Leibold et al. 2004; Logue et al. 2011).

There are currently at least three different lines of research towards such a unified description. The first builds on patch occupancy models, which simulate the stochasticity of patch occupancy, a consequence of demographic stochasticity, based on explicit colonisation–extinction probabilities (Ovaskainen 2002; Moilanen 2004). Recent variants consider direct time-dependence of these probabilities (Bertassello et al. 2021) or a dependence on the occupation of patches by other species (Jackson et al. 2025). Amongst the limitations of these approaches are difficulties in describing complex indirect interactions between species, which can lead to species sorting through competitive exclusion, and of mass effects that can override such exclusion. The alternative are models describing metacommunities through systems of coupled ordinary differential equations (ODEs; e.g., Thompson et al. 2017; O’Sullivan et al. 2019, 2021), which intrinsically represent species sorting and mass effect but struggle (as we demonstrate below) capturing the discrete nature of immigration and extirpation events. A third approach, the stochastic metacommunity models proposed by Lerch et al. (2023), provide detailed descriptions of the joint probability distributions of populations sizes resulting from stochastic population dynamics within patches, dispersal, and patch occupancy dynamics, but require fully connected networks of identical patches to remain computationally feasible.

Here we introduce the probabilistic–stochastic–deterministic (PSD) approximation, a framework that bridges computationally expensive IBMs and analytically tractable but incomplete ODE models. The key innovation is a hybrid state representation. Species populations at each patch exist in one of three states requiring different kinds of descriptions: largely excluded due to negative local invasion growth rate, permitting a probabilistic description; ‘waiting’, because their invasion growth rate is positive but they have not established, yet, modelled stochastically; and established, permitting deterministic ODE modelling. Transitions between states capture colonisation and extirpation explicitly. This structure naturally encompasses all four paradigms: patch dynamics emerges from stochastic state transitions; species sorting from spatially varying growth rates and biotic interactions; mass effects when high dispersal rates and small body size reduce PSD to ODE dynamics; and demographic stochasticity through the probabilistic establishment process.

Missing in our current PSD formulation is only a describe of extirpations of small established populations through demographic stochasticity (Kessler and Shnerb 2015), but an extension to include this process might be possible. We do not consider this a crucial omission, as such stochastic extirpation can occur only with very small population sizes. In general, seemingly random disappearances of local populations are more naturally explained as resulting from diffuse competitive or antagonistic interactions in complex ecological networks (O’Sullivan et al. 2023a,b; Nwankwo and Rossberg 2026; Cockrell et al. 2024), a phenomenon fully captured by the PSD approximation. After introducing our modelling framework, we first demonstrate how ODE models fail at low invasion rates and show that PSD accurately reproduces IBM dynamics across the full parameter space. We then demonstrate two applications using rock–paper–scissors metacommunities with intransitive competition (Kerr et al. 2002; Reichenbach et al. 2007): spatial pattern formation, and analytical derivation of slow collective metacommunity oscillations under long-distance dispersal, a result inaccessible to both ODE models (lacking stochasticity) and IBMs (lacking analytical tractability). Because PSD avoids simulating the many uninformative demographic fluctuations that IBMs must track, and can employ larger integration time steps without loss of accuracy, PSD is potentially much faster than IBM, particularly for systems with large established populations. For example, in our single-patch validation simulations with *S* = 300 species over *T* = 10,000 time units, PSD completed in 1.1 s versus 2.0 s for IBM on a single CPU core, a factor of 1.8 speedup arising because PSD’s smooth expected-value dynamics permit time steps of Δ*t* = 10 whereas IBM requires Δ*t* = 1 to resolve individual-level demographic events.

## 2 Models

We compare three modelling approaches of increasing mechanistic detail: ODE models, IBMs, and our PSD approximation. All three frameworks share a common ecological foundation but differ in their treatment of demographic stochasticity.

### 2.1 Ordinary differential equation (ODE) model

To illustrate our approach, we initially consider community dynamics at a single patch. The time-dependent state is described by the vector **B**(*t*) = (*B*_1_(*t*), …, *B*_*S*_(*t*)) of biomass abundances of *S* species, with dynamics

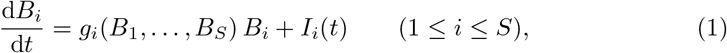

where *g*_*i*_(·) is the instantaneous per-capita growth rate and *I*_*i*_(*t*) ≥ 0 is the immigration flux. In our examples, we employ a competitive generalised Lotka–Volterra model (Case 2000) with

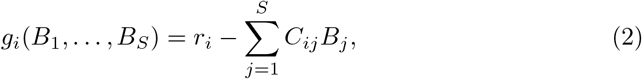

where *r*_*i*_ are the intrinsic growth rates and the coefficients *C*_*ij*_ quantify competitive effects. In our simulations, we set *r*_*i*_ = *C*_*ii*_ = 1 for all species, thus normalising biomass such that single-species carrying capacity equals unity. In principle, variation of intrinsic growth rates between species and patches could represent environmental filtering, but we keep *r*_*i*_ constant here to isolate demographic effects. Unless otherwise noted, off-diagonal competition coefficients are randomly set to 0.4 with probability 0.4 and otherwise to zero (Rossberg 2013; O’Sullivan et al. 2019).

For metacommunity simulations, we extend Eq. (1) to a spatial grid of *L*_*x*_ × *L*_*y*_ patches, with dispersal fluxes *I*_*i,x,y*_(*t*) = ∑_(*x′*, *y′*) ≠ (*x,y*)_ *K*_(*x,y*),(*x′*,*y′*)_ *B*_*i,x′,y′*_ (*t*), where *K* is a fixed dispersal kernel. Numerical implementation details are provided in Supplementary Information S1.

### 2.2 Individual-based model

The IBM provides a stochastic reference for validation. It tracks the abundances *N*_*i*_(*t*) of discrete individuals of clonally reproducing species *i*, thus representing a minimal model capturing demographic stochasticity. The corresponding biomasses are *B*_*i*_(*t*) = *N*_*i*_(*t*) · *m*_*i*_, where *m*_*i*_ is the adult body mass of species *i* (as common, we skip over pre-adult stages for simplicity). When all *m*_*i*_ are small, the law of large numbers ensures IBM dynamics approximate the ODE; when some *m*_*i*_ are large, demographic stochasticity becomes relevant. In our numerical examples, we set all *m*_*i*_ to the same value *m*_0_.

The model tracks three elementary events (Table 2). Birth and mortality rates are constructed such that their difference equals net local biomass change *g*_*i*_*N*_*i*_*m*_*i*_ from the ODE. We keep mortality fixed at *M*_*i*_ as much as possible while ensuring both rates remain non-negative (Black and McKane 2012). Our simulation algorithm, with discrete time steps, is detailed in Supplementary Information S2. In numerical examples we choose the same value *M* = 0.2 for all *M*_*i*_.

**Table 1:** Coverage of the four metacommunity paradigms by different modelling approaches. PD: Patch dynamics (colonisation and extinction); SS: Species sorting (environmental filtering and biotic interactions); ME: Mass effects (deterministic dispersal influence); DS: Demographic stochasticity. A check mark (✓) denotes explicit representation; a circle (◦) denotes non-coverage, a tilde ~ partial coverage.

| Modelling approach | PD | SS | ME | DS |
| --- | --- | --- | --- | --- |
| Individual based models | ✓ | ✓ | ✓ | ✓ |
| Stochastic metacommunity models | ✓ | ✓ | ✓ | ✓ |
| Stochastic patch occupancy | ✓ | ○ | ○ | ✓ |
| ODE models | ~ | ✓ | ✓ | ○ |
| **This work (PSD framework)** | ✓ | ✓ | ✓ | ~ |

**Table 2:** Elementary events in the stochastic individual-based model. Each event type, its effect on individual count *N*_*i*_, and the rate at which it occurs. The net growth rate *g*_*i*_ = *r*_*i*_ − ∑_*j*_ *C*_*ij*_*B*_*j*_ can be positive or negative; *M*_*i*_ denotes the baseline mortality rate of species *i* and *I*_*i*_ its immigration flux.

| Event | Transition | Rate per patch |
| --- | --- | --- |
| Birth | $N_i \rightarrow N_i + 1$ | $\max(0, g_i + M_i) \cdot N_i$ |
| Death | $N_i \rightarrow N_i - 1$ | $\max(-g_i, M_i) \cdot N_i$ |
| Immigration | $N_i \rightarrow N_i + 1$ | $I_i/m_i$ |

### 2.3 Probabilistic–stochastic–deterministic (PSD) approximation

The PSD approximation assigns each species population to one of three states described by different mathematical approaches. Our specific formulation relies on the IBM assumption detailed in Table 2 that, for growth rates *g*_*i*_ near zero, mortality is constant *M*_*i*_ while the birth rate *g*_*i*_ + *M*_*i*_ is density dependent through *g*_*i*_. This is ecologically plausible: offspring generation is often resource-limited, while adult survival depends more strongly on phenotype.

#### 2.3.1 State characterisation

The distinction between states depends on the sign of the *non-self growth rate*

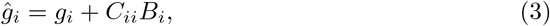

which generalises invasion growth rate to established populations such that it is generally positive for established species. This approach is valid when the equilibrium biomasses of established species tend to substantially exceed expected biomasses of non-established species, as in Fig. 2.

PSD classified populations with *ĝ*_*i*_ < 0 as being in the P-state (probabilistic), described by the expectation value of biomass, which is here also denoted by *B*_*i*_ and evolves according to Eq. (1) (Kendall 1948). Patches with *B*_*i*_ ≪ *m*_*i*_ are unoccupied with high probability.

Populations with *ĝ*_*i*_ > 0 are either in the S-state (stochastic) if not yet established or the D-state (deterministic) if established. In the S state, dynamics are described by two variables. The variable *B*_*i*_ represents the expected biomass of propagules that have arrived but fail to establish. Its dynamics are approximated by

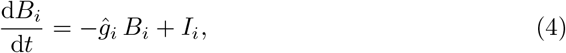

where the negative sign arises because we track only failed establishment attempts. The second variable is a *Poisson clock P*_*i*_, which reaches zero from below when an establishing propagule arrives. The randomness of immigration is modelled by initialising *P*_*i*_ = ln(*U*) with *U* randomly sampled from]0; 1[, and dynamics are given by

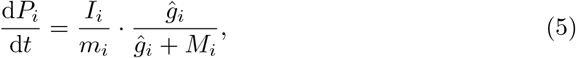

where the two factors represent propagule arrival rate and establishment probability, respectively. In the D state, biomass follows Eq. (1), with demographic stochasticity disregarded. When evaluating intrinsic growth rates *g*_*i*_, as in Eq. (2), we approximate the small biomasses of species in P- and S-states by their expectation values *B*_*i*_. Derivations are provided in Supplementary Information S3.

#### 2.3.2 Transition dynamics

Transitions from D-to P-states as *ĝ*_*i*_ becomes negative are silent as both follow Eq. (1). When *ĝ*_*i*_ becomes positive, a P-state population transitions to either D or S depending on propagule presence and chance. Building on analytic theory for birth-death processes (Branson 1991), we approximate the probability of transitioning to S as

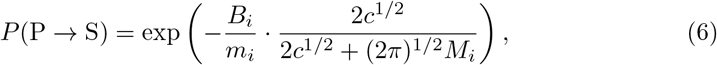

where *c* = *dĝ*_*i*_*/dt >* 0 is the rate at which *ĝ*_*i*_ crosses zero. Propagules present when *ĝ*_*i*_ becomes positive typically fail to establish if this transition is slow (*c*^1*/*2^ ≪ *M*_*i*_). Populations in S state transition to D when *P*_*i*_ reaches zero (successful establishment) or to P if *ĝ*_*i*_ becomes negative.

To maintain consistency with expectation-value dynamics, biomass is adjusted at transitions to D state: at transitions from P, we replace the expected biomass *B*_*i*_ by *B*_*i*_*/*(1 − *P* (P → S)); for transitions from S we set D-state biomass to *m*_*i*_(*ĝ*_*i*_ + *M*_*i*_)*/ĝ*_*i*_ (details in Supplementary Information S3).

## 3 Single-patch validation: comparing ODE, IBM, and PSD

We compared the three modelling frameworks to establish if PSD successfully captures IBM dynamics where ODE fails. We simulated communities of *S* = 300 species, with immigration flux *I*_*i*_ = 10^−8^ per species. To explore effects of demographic stochasticity, we compared two body-mass regimes: low body-mass (*m*_0_ = 10^−11^, many small individuals) and high body-mass (*m*_0_ = 10^−4^, few large individuals). All other parameters were given above. Each simulation ran for *T* = 10^4^ unit times.

### 3.1 Biomass dynamics and distributions

Figure 1 presents biomass time series under both regimes. At low body-mass (left column), all three models generate visually indistinguishable vigorous population fluctuations, known similarly from previous studies (Mallmin et al. 2024). The similarity of the three model outputs reflects the law of large numbers (Kurtz 1970): demographic stochasticity averages out when populations comprise many small individuals. Occasional brief pauses in turnover dynamics, visible for example around *t* = 9200 in IBM and *t* = 9500 in PSD, occur with all three model types and reflect the finite species pool.

**Fig. 1:**
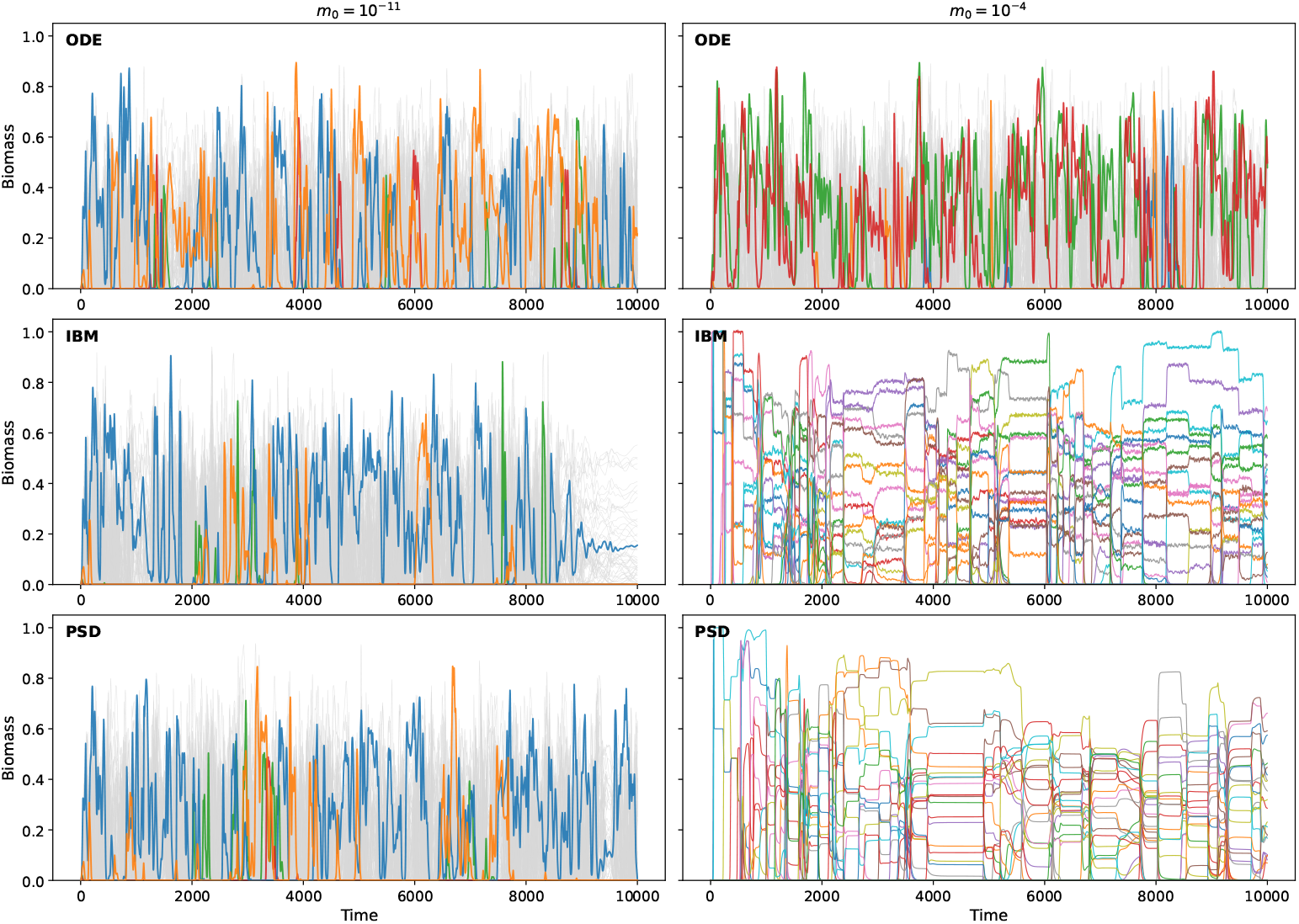
Biomass trajectories under two body-mass regimes. Left: *m*_0_ = 10^−11^; right: *m*_0_ = 10^−4^. Top: ODE; middle: IBM; bottom: PSD. At low *m*_0_, all models agree. At high *m*_0_, ODE shows misleading smooth dynamics; IBM and PSD correctly capture discrete colonisation–extinction events.

At high body-mass (right column), model outputs differ. ODE model dynamics remain unchanged, as they do not depend on body mass. However, with just *I*_*i*_*/m*_0_ = 10^−6^ individuals arriving per time unit and species, colonisation is actually rare and discrete, a pattern captured only by IBM and PSD. The IBM shows abrupt colonisation–extinction events with demographic fluctuations around intermittent equilibria; PSD reproduces these episodic dynamics through its Poisson arrival process (Eq. 5) and establishment probability (Eq. 6), but follows deterministic trajectories between these transitions.

Figure 2 shows log-biomass distributions sampled from the stationary regime after discarding an initial 70% burn-in period. At low body-mass (Fig. 2a), all three models produce nearly identical bimodal distributions. The bimodal structure, arising for small immigration fluxes *I*_*i*_, was explained in analyses by Arnoulx de Pirey and Bunin (2024) and Mallmin et al. (2024). Essentially, the lower peak corresponds to the equilibrium biomass

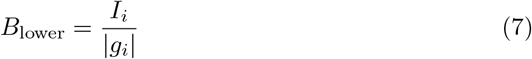

balancing typical negative values of *g*_*i*_ with the immigration flux *I*_*i*_, and the upper peak corresponds, for Lotka–Volterra competition models of the form of Eq. (2), to the equilibrium biomass

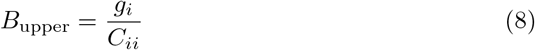

for typical positive values of *g*_*i*_. Intermediate biomasses arise because *g*_*i*_ varies randomly through time between positive and negative values.

**Fig. 2:**
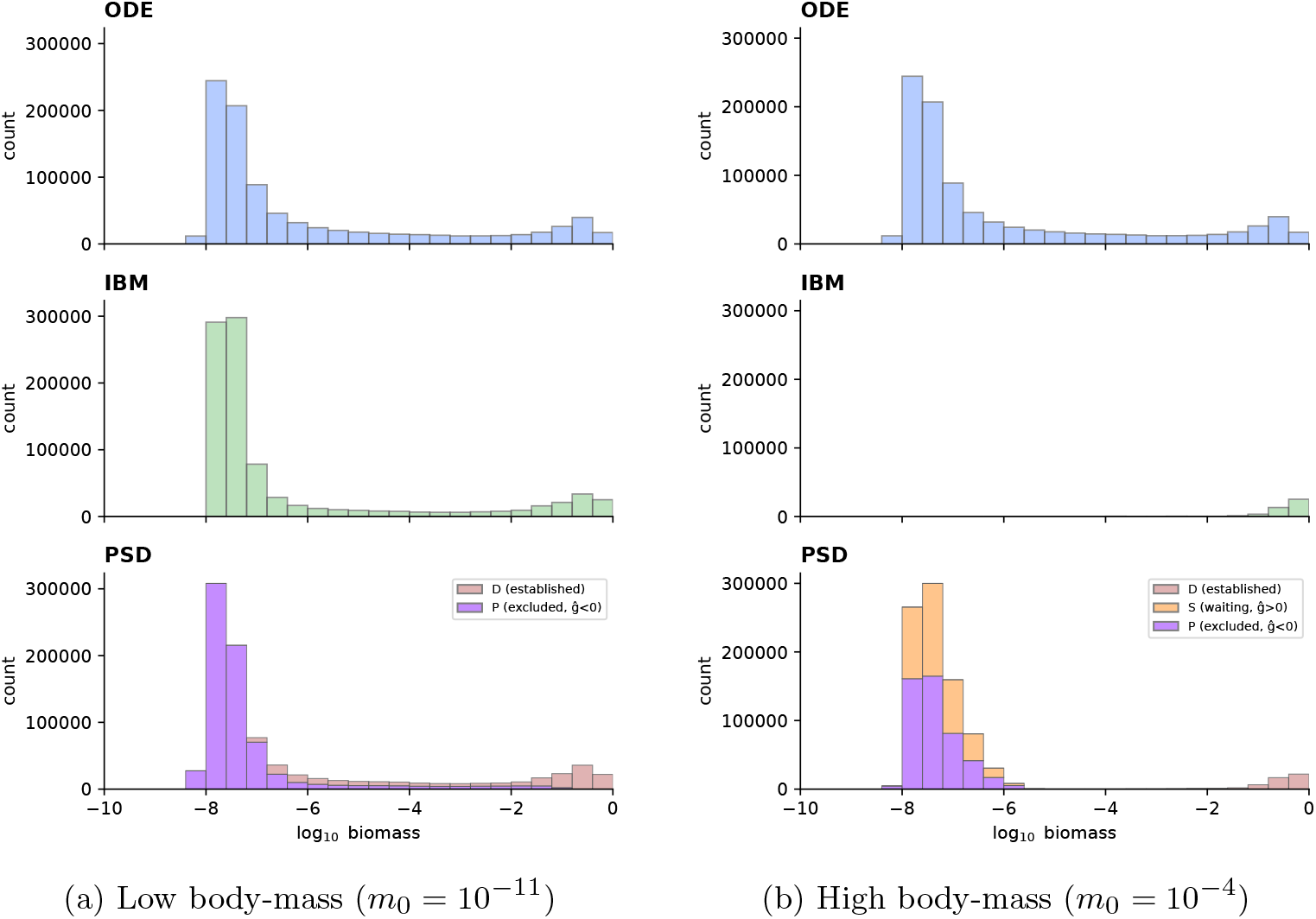
Biomass distributions in the stationary regime. Histograms of log_10_ *B*_*i*_ for all species, sampled after a 70% burn-in: ODE (top), IBM (middle), PSD (bottom). For PSD, colours indicate state: D (established, red), S (waiting, orange), P (excluded, purple). (a) Low body-mass: all models produce identical bimodal distributions. Because *m*_0_ is very small, transitions into S-state are rare (Eq. 6); species flip directly between D (*ĝ* > 0) and P (*ĝ* < 0) states, yielding roughly equal D/P proportions at intermediate biomass. (b) High body-mass: ODE remains bimodal and unchanged; IBM shows unimodal distribution near carrying capacity; PSD separates D-state (established, *B >* 0.01) from S-state (waiting, *ĝ*_*i*_ > 0).

We can complement the analyses of Arnoulx de Pirey and Bunin (2024) and Mallmin et al. (2024) with an approximate analytic theory for the distribution of *g*_*i*_ by Cockrell et al. (2024) for fixed *r*_*i*_ = *r* and *C*_*ii*_ = 1 and off-diagonal elements *C*_*ij*_ sampled independently from a given distribution with mean 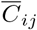 and variance var *C*_*ij*_. According to this theory, *g*_*i*_ is zero on average and has a standard deviation

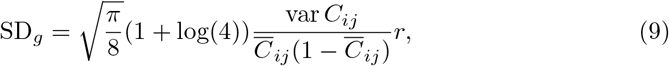

where the numerical prefactor evaluates to 1.495. By representing typical positive and negative values of *g*_*i*_ by *±*SD_*g*_, we obtain for our example with *I*_*i*_ = 10^−8^ the values log_10_ *B*_lower_ = −7.6 and log_10_ *B*_upper_ = −0.4, in reasonably good agreement with Fig. 2a.

At high body-mass (Fig. 2b), distributions differ: the ODE result is independent of body-mass and so remains unchanged, while IBM and the D-states of PSD are unimodally distributed with population biomasses close to unity.

For PSD, we additionally plot histograms for the small expected biomasses in S- and P-states, which have a mode at a position similar to the lower mode of the ODE model.

In simple words, the ODE fails to reproduce the individual-based dynamics with large body mass because the corresponding arrival rate of individuals (*I*_*i*_*/m*_0_ = 10^−6^) is small, while all other dynamic parameters are of order unity. The ODE wrongly describes ‘fractions of individuals’ arriving during a unit time, which then either shrink further (if *ĝ*_*i*_ < 0) or grow (if *ĝ*_*i*_ > 0) to first become complete individuals and then entire populations. In the underlying IBM, none of this happens. Instead, entire individuals arrive occasionally and some of these establish new local populations (Fig. 1, *m*_0_ = 10^−4^).

### 3.2 Statistical comparison

Table 3 presents species richness (number of species with *B*_*i*_ > 10^−3^) and invasion rates, computed as means over the stationary time series after burn-in. We tested PSD across three body-mass regimes: low (*m*_0_ = 10^−11^), intermediate (*m*_0_ = 10^−7.6^, where *m*_0_ ≈ *B*_lower_), and high (*m*_0_ = 10^−4^).

**Table 3:** Summary statistics for single-patch simulations. Mean *±* SE across *n* = 10 time-averaged chunks.

| Regime | Model | Richness | Invasion rate |
| --- | --- | --- | --- |
| Low $m_0$ ($10^{-11}$) | ODE | $47.04 \pm 0.38$ | $(3.39 \pm 0.04) \times 10^{-2}$ |
| | IBM | $46.43 \pm 0.18$ | $(3.37 \pm 0.06) \times 10^{-2}$ |
| | PSD | $47.51 \pm 0.40$ | $(3.48 \pm 0.05) \times 10^{-2}$ |
| Intermediate $m_0$ ($10^{-7.6}$) | ODE | $48.84 \pm 1.10$ | $(3.87 \pm 0.26) \times 10^{-4}$ |
| | IBM | $50.86 \pm 0.86$ | $(4.39 \pm 0.20) \times 10^{-4}$ |
| | PSD | $48.91 \pm 0.66$ | $(3.70 \pm 0.21) \times 10^{-4}$ |
| High $m_0$ ($10^{-4}$) | ODE | $48.05 \pm 0.18$ | $(3.52 \pm 0.02) \times 10^{-2}$ |
| | IBM | $18.09 \pm 0.93$ | $(8.25 \pm 1.29) \times 10^{-5}$ |
| | PSD | $15.06 \pm 0.39$ | $(1.26 \pm 0.09) \times 10^{-4}$ |

At low body mass, ANOVA revealed no significant differences among models (*F*_2,27_ = 0.29, *p* = 0.75). At intermediate body-mass, where the ratio *B*_lower_*/m*_0_ ≈ 1 presents the most challenging test for the PSD approximation, models again showed no significant differences (*F*_2,27_ = 1.65, *p* = 0.21); IBM and PSD remained statistically indistinguishable (Tukey HSD: *p* = 0.29). At high body-mass, differences were highly significant (*F*_2,27_ = 3.17 × 10^3^, *p <* 10^−27^): ODE predicts approximately threefold higher richness than IBM or PSD. Critically, IBM and PSD are statistically indistinguishable across all three regimes (Tukey HSD: *p* = 0.86 at high *m*_0_), confirming that PSD captures IBM dynamics over the entire body-mass range.

These results validate PSD as an efficient alternative to IBM in the entire body-mass range from weak demographic stochasticity (low *m*_0_) through the critical transition zone where *m*_0_ ≈ *B*_lower_ (intermediate *m*_0_) to strong demographic stochasticity (high *m*_0_).

## 4 Multi-patch metacommunity dynamics

After this single-patch validation, we turn to spatially explicit metacommunities using the classic May-Leonard (May and Leonard 1975) rock–paper–scissors (RPS) system (Kerr et al. 2002; Reichenbach et al. 2007), a minimal model exhibiting cyclic competition and spatial pattern formation that tests whether PSD captures emergent spatio-temporal dynamics.

### 4.1 Rock–paper–scissors metacommunity

We consider *S* = 3 species with cyclic competition on an *L*_*x*_ × *L*_*y*_ grid with periodic boundaries. The competition matrix is

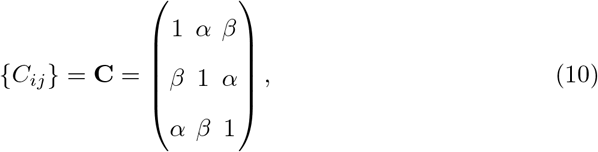

where *α* = 1.7 and *β* = 0.4. Species 1 suppresses species 3, species 2 suppresses species 1, and species 3 suppresses species 2.

### 4.2 Spatial pattern formation

In the first example, we use nearest-neighbour dispersal on a 50 × 50 grid, where each patch exchanges propagules with its four von-Neumann neighbours at rate *D* = 2 × 10^−7^ (Supplementary Information S6). Figure 3 presents snapshots of spatial species distributions across the three modelling frameworks. The ODE (Fig. 3A) produces only fine-grained spatial structure. As in the single-patch case, immigrations in the ODE are too frequent because arbitrarily small biomass fluxes translate into continuous propagule arrival. Therefore domains dominated by a single species remain too small to form coherent spatial patterns. The IBM (Fig. 3B) generates large coherent domains dominated by single species that evolve through cyclic succession (red → blue → green → red), creating dynamic mosaics characteristic of RPS systems (Reichenbach et al. 2007; Kerr et al. 2002). The PSD (Fig. 3C) reproduces IBM’s qualitative features: coherent patches emerge, separated by fronts propagating through the metacommunity (last four snapshots in Fig. 3B,C). With IBM, sites change dominant species at an average rate of 1.88 × 10^−4^ per time unit and similarly at 1.97 × 10^−4^ for PSD, while for ODE this rate is 4.86 × 10^−3^, roughly 25-fold higher. This confirms that PSD captures metacommunity-scale stochastic dynamics at high accuracy but a fraction of the computational cost.

**Fig. 3:**
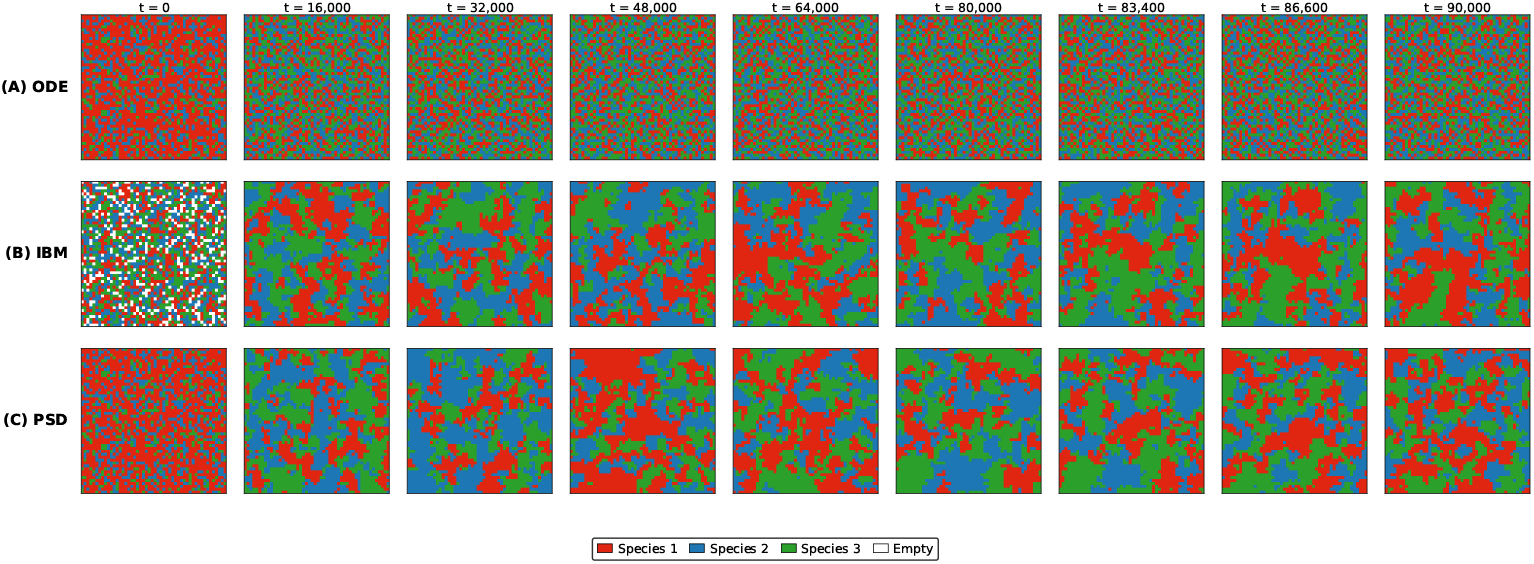
Spatial pattern formation in the RPS metacommunity. Snapshots at *t* = 0, 16,000, 32,000, 48,000, 64,000, 80,000, 83,400, 86,600, 90,000. Colours: species 1 (red), species 2 (blue), species 3 (green); white = empty. **(A)** ODE: fine-grained structure; immigration is too frequent for coherent patches to form. **(B)** IBM: large coherent patches with cyclic succession. **(C)** PSD: accurately reproduces IBM spatial dynamics at lower computational cost.

These results demonstrate how the ODE model fails to capture spatial pattern formation driven by discrete demographic events. It is likely that ODE models would similarly fail to reproduce shifting-mosaic structures observed in real landscapes (Wang and Finley 2011). By contrast, PSD captures the essential IBM dynamics.

## 5 Analytical prediction of metacommunity oscillation periods

The preceding sections established that PSD accurately reproduces IBM dynamics. We now demonstrate that the PSD framework enables analytical predictions inaccessible to either ODE or IBM alone.

### 5.1 Derivation of the oscillation period

We consider the limit of global (long-distance) dispersal, where propagules leaving a patch at total emigration rate *D* per individual and reach any other patch with equal probability. In the RPS system, species invade patches dominated by their inferior competitor at a rate given by the product of dispersal flux, establishment probability, and patch availability, all quantities represented explicitly in the PSD approximation. Linearising the resulting mean-field dynamics for patch occupancy around the symmetric equilibrium 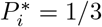 and computing eigenvalues (Supplementary Information S7), we find metacommunity occupancy to oscillate with period

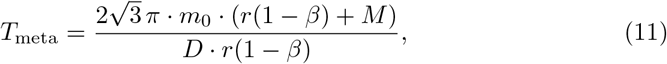

independent of *α*, where *α* and *β* are the dominant and subdominant competition coefficients from Eq. (10). The period scales linearly with body mass *m*_0_ and inversely with dispersal rate *D*.

Remarkably, the non-linear mean-field equations for this process take the mathematical form of a three-species Lotka–Volterra model with antisymmetric competition, for which the oscillation amplitude is known to be a conserved quantity (Volterra 1931). Owing to the fixed number of available patches, this structure and the resulting conservation law remain intact even when vital rates and competition coefficients vary between species. In simulations and in nature we correspondingly expect oscillations with indeterminate, slowly varying amplitudes.

### 5.2 Contrast with ODE predictions

The metacommunity period (Eq. 11) differs fundamentally from isolated-patch ODE dynamics. For a single patch, linearization around the coexistence equilibrium yields (May and Leonard 1975; Hofbauer and Sigmund 1998)

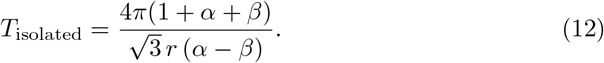

Crucially, *T*_isolated_ is independent of body mass *m*_0_ and dispersal rate *D*. For our RPS parameters (Section 4), we obtain *T*_isolated_ ≈ 17 unit times and *T*_meta_ ≈ 2902 unit times, illustrating the difference between these two kinds of oscillations.

### 5.3 Numerical validation

Figure 4 validates these predictions. At full dispersal rate *D* = 5 × 10^−7^, both IBM and PSD exhibit oscillations with period ≈ 2900 unit times, matching Eq. 11. When dispersal is halved to *D* = 2.5 × 10^−7^, the observed period doubles to ≈ 5800 exactly as predicted. The ODE fails entirely to reproduce these slow cycles, instead showing rapid boundary dynamics determined by *T*_isolated_, first in clusters, then synchronised over all patches, slowing down as the heteroclinic cycle linking the three single-species equilibria is approached (Hofbauer 1994).

**Fig. 4:**
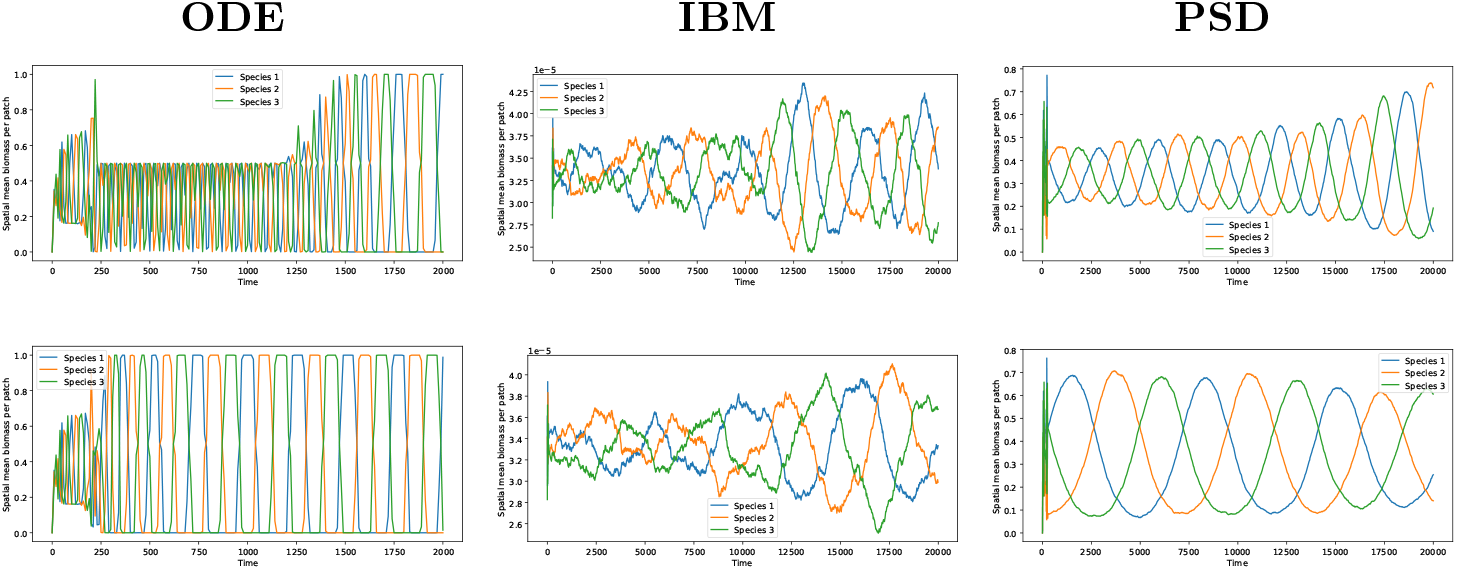
Slow metacommunity oscillations under long-distance dispersal. Mean biomass per patch for RPS species. **Top:** *D* = 5 × 10^−7^. **Bottom:** *D* = 2.5 × 10^−7^. ODE (left) shows rapid boundary dynamics. IBM (centre) and PSD (right) exhibit slow oscillations matching Eq. 11. Halving *D* doubles the period, confirming the metacommunity mechanism.

## 6 Discussion

### 6.1 Innovation and limitations

Above, we introduced the Probabilistic–Stochastic–Deterministic (PSD) approximation of metacommunity dynamics and demonstrated its good agreement with the ground-truth represented by an IBM, and also with a corresponding ODE model for parameters where the ODE model is valid. Our theory remains incomplete insofar as it does not capture extirpation of established populations through demographic stochasticity, implying a condition that *m*_0_ ≪ *B*_upper_. We speculate that this could be captured in a model variant running Poisson clocks for D-state populations tuned to capture the likelihood of such extirpations, e.g., based on Leigh (1981). The condition *m*_*i*_ ≪ *B*_upper_ also justifies our approximation of *g*_*i*_ by *ĝ*_*i*_. The good agreement that we find between IBM and PSD simulations regardless of these simplifying assumptions demonstrates that, while demographic stochasticity undoubtedly plays a role in the final phase of a species’ extirpation, the associated randomness is not generally relevant for higher-level community and metacommunity dynamics.

Our current formulation is also restricted to the case where the density dependence of demographic rates is in the birth rates only. An extension to density dependence in both birth and death rates appears feasible but would require further work. Other limitations of our theory in the current formulation include the absence of stage structure and of sexual reproduction (both of which can affect dispersal processes). We encourage research to uncover the implications of these limitations for metacommunity ecology.

### 6.2 Unification of metacommunity paradigms

To explain how the PDF approximation describes the four classical metacommunity paradigms, we distinguish between, on the one hand, the key phenomena underlying each paradigm (mass effects, patch dynamics, species sorting, and demographic stochasticity) on and, on the other hand, idealised scenarios where the paradigms arise in ‘pure’ form.

#### 6.2.1 Uniform descriptions of key phenomena

Mass effects and patch dynamics arise depending on the relative values of *B*_lower_, given by Eq. (7) above, and *m*_*i*_. In Fig. 1 and 2, we illustrated that deterministic descriptions of mass effects using ODEs remain valid when the theoretical equilibrium population biomass for typical mass effects, *B*_lower_, is much larger than *m*_*i*_. In this case, robust sink populations (in P-state) can establish that will predictably grow into established (D-state) populations once *g*_*i*_ becomes positive. In the opposite case, *m*_*i*_ ≪ *B*_lower_, immigrations are too rare for sink populations to establish. No mass effects arise. In this case, transitions between P- and D-state, by Eq. (6), go through the S-state, resulting in patch dynamics. The critical role of the dimensionless parameter *B*_lower_*/m*_*i*_ in this context becomes plausible when noting that it corresponds to the ratio between the rate parameters for arrival of individuals (*I*_*i*_*/m*_*i*_) and for typical population decline |*g*_*i*_|.

Since our approximation requires *m*_*i*_ ≪ *B*_upper_, there are three different kinds of orderings left to consider for the magnitudes of *B*_lower_, *B*_upper_, and *m*_*i*_. These are *B*_lower_ ≪ *m*_*i*_ ≪ *B*_upper_ (patch dynamics), *m*_*i*_ ≲ *B*_lower_ ≪ *B*_upper_ (mass effects), and the exotic case where *m*_*i*_ ≪ *B*_upper_ ≲ *B*_lower_ (with *B*_lower_ and *B*_upper_ given by Eqs. (7) and (8) above). We shall briefly discuss the last case for completeness. By Eqs. (7) and (8), it implies that, by order of magnitude, |*g*_*i*_|^2^ ≲ *C*_*ii*_*I*_*i*_. This situation can arises, e.g., in farmland, where, due to intense sowing (large *I*_*i*_), populations establish with *B*_*i*_ ≈ (*I*_*i*_*/C*_*ii*_)^1*/*2^ ≈ *B*_upper_ (*B*_lower_*/B*_upper_)^1*/*2^ ≫ *m*_*i*_ regardless of the local viability (*g*_*i*_) of species *i*. The PSD formalism predicts for this cases that transitions into the S-state are rare by Eq. (6) (since *B*_*i*_ ≫ *m*_*i*_) and that, where they occur, they are short lived by Eq. (5): the typical duration of S-states *m*_*i*_*/I*_*i*_ is small compared to the characteristic time scale (*C*_*ii*_*I*_*i*_)^−1*/*2^ because *m*_*i*_*/I*_*i*_ ≪ *B*_upper_*/I*_*i*_ = |*g*_*i*_|*/*(*C*_*ii*_*I*_*i*_) ≲ |*g*_*i*_|*/*(*C*_*ii*_*I*_*i*_) × [*C*_*ii*_*I*_*i*_*/*|*g*_*i*_|^2^]^1*/*2^ = (*C*_*ii*_*I*_*i*_)^−1*/*2^. Hence, the PSD approximation reduces to an ODE model in this case, just as for mass effects.

Species sorting is accurately described by the PSD approximation through dependencies of *g*_*i*_ on local environment and local community. Specifically, the momentary growth rates *g*_*i*_ take, through their dependence on *B*_1_, …, *B*_*S*_, account of the effects of all established populations (in D-state), and, in expectation (up to non-linear corrections), of the small effects of all non-established populations (in P-state and S-state). In case of the Lotka–Volterra form for *g*_*i*_, Eq. (2), abiotic (environmental) filtering results from variation of the *r*_*i*_ between species and patches, while biotic filtering results from the sum over interaction terms.

Demographic stochasticity is captured by the PSD approximation where it matters: in (i) the random arrival of propagules at patches (first factor in Eq. (5)), (ii) the establishment probability of propagules (second factor Eq. (5)), (iii) the establishment of sink populations when invasion growth rate turns positive (Eq. (6)).

#### 6.2.2 Paradigmatic metacommunity scenarios

Building on above considerations, metacommunity models with pure mass effects arise in metacommunities in the limit *m*_0_ → 0 with fixed dispersal rates.

To construct a scenario where pure patch dynamical models become valid, dispersal rates need to be sufficiently low such that *B*_lower_ ≪ *m*_0_ in all circumstances. Patches will then spend most time in a local equilibrium (or other attractor) given by their D-state community. This equilibrium is only occasionally interrupted by random immigrations that disturb the community, often extirpate residents, and result in a new D-state community (Cockrell et al. 2024). These immigrations and the extirpations they trigger become essentially random from the point of view of a single species (such that patch-dynamical metapopulation models apply) when the compositions of the D-state communities differs substantially between the patches amongst which a species disperses (high spatial beta diversity) and also changes through time (high temporal beta diversity) (O’Sullivan et al. 2021; Cockrell et al. 2024). Such a situation naturally arises with long-range dispersal when regional species pool (gamma diversity) far exceeds the intrinsic limit to local (alpha) diversity imposed by ecological structural instability (O’Sullivan et al. 2019).

Pure species sorting arises with low dispersal rates in the opposite extreme where regional species pools are so small that a static equilibrium arises in which all populations are either in D- or in P-states (no S-state). By the theory of Bunin (2017), this requires gamma diversity to remain below around twice the upper bound to alpha diversity.

The PSD approximation cannot recover neutral theory as originally conceived as a limiting case, because neutral population dynamics correspond to the situation where *g*_*i*_ is always zero, a singular case in the PSD approximation. However, the mathematics of neutral theory can be re-interpret as referring, rather than to the numerical sizes of species populations, to the numbers of patches occupied by species (O’Sullivan et al. 2023a), thus describing “neutral dispersal processes” sensu Cottenie (2005). We expect that models approximated by this patch-level neutral theory arise in the patch-dynamical regime of the PSD-approximation when all species are statistically equivalent (same distribution of interaction strengths *C*_*ij*_) all patches ecologically equivalent for all species (all *r*_*i*_ identical).

### 6.3 Emergent concepts

As a general principle, one and the same complex system can often be described by a variety of different models that resolve the trade-off between simplicity of model formulation, computation speed, and accuracy in different ways (Rossberg 2007)

Often one finds that more elaborate but computationally more efficient model formulations introduce new concepts, which, on this basis, can be pin-pointed as concepts relating to emergent high-level phenomena. Usually, these emergent phenomena are empirically known before being used in models. However, the modelling helps sharpening definitions and understanding the emergent phenomena in context.

In the case of the PSD approximation, we obain primarily the conceptual distinction between species populations in P-, S-, and D-states. The concept of D-state populations is readily identified with the ecological concept of populations resident at a patch, offering a sharp mathematical distinction from vagrants or transient individuals and their offspring, represented in our formulation by P- and S-state populations.

Competitively excluded populations directly correspond to P-states. As discussed above, these can become sink populations at high dispersal rates. The concept of dark diversity, “ecologically suitable species that are absent from a site but present in the surrounding region” (Pärtel et al. 2025) is related to S-state populations.

### 6.4 Methodological resolution

The PSD formalism resolves a long-standing methodological tension between stochastic individual-based simulations and deterministic approximations. Deterministic approaches, from reaction-diffusion partial differential equation models to matrix metapopulation models, offer analytical insights into spatial spread and persistence (Hastings and Botsford 2006), but fundamentally fail to capture demographic stochasticity governing, e.g., colonisation at invasion fronts. Individual-based models correctly handle discrete stochastic events (Bocedi et al. 2014) but sacrifice mathematical transparency and computational efficiency.

PSD bridges this divide by mathematically partitioning the colonisation process: the biomass variables in P- and S-states follow tractable linear dynamics describing dispersal across landscapes, while the Poisson clocks of S-states describe the stochasticity of establishment. This retains the predictive power of analytical models for transport while incorporating the demographic realism of IBMs.

## 7 Conclusion

We developed the Probabilistic–Stochastic–Deterministic (PSD) approximation as a unified framework for metacommunity dynamics. The PSD approximation accurately reproduces IBM dynamics while running in some potentially orders of magnitude faster, and enables analytical theory, such as for the metacommunity oscillation period, inaccessible to either ODE models (lacking stochasticity) or IBM (lacking tractability) alone. Deterministic ODE metacommunity models are valid only as long as arrival rates of individuals at patches are large compared to typical rates of population decline or growth.

The four paradigms of metacommunity theory are not in conflict but represent valid descriptions in different regions of parameter space, which we have identified. This resolves a long-standing tension in metacommunity ecology: apparent conflicts between paradigms arose because different studies implicitly operated in different parameter regimes. The PSD framework provides a unified description encompassing all regimes and, importantly, the large swaths of parameter space in between.

## References

Bertassello LE, Bertuzzo E, Botter G, et al (2021) Dynamic spatio-temporal patterns of metapopulation occupancy in patchy habitats. Royal Society Open Science 8(1):201309. 10.1098/rsos.201309

Black AJ, McKane AJ (2012) Stochastic formulation of ecological models and their applications. Trends in Ecology & Evolution 27(6):337–345. 10.1016/j.tree.2012.01.014

Bocedi G, Palmer SCF, Pe’er G, et al (2014) RangeShifter: a platform for modelling spatial eco-evolutionary dynamics and species’ responses to environmental changes. Methods in Ecology and Evolution 5(4):388–396. 10.1111/2041-210X.12162

Branson D (1991) Inhomogeneous birth-death and birth-death-immigration processes and the logarithmic series distribution. Stochastic Processes and their Applications 39(1):131–137. 10.1016/0304-4149(91)90037-D

Bunin G (2017) Ecological communities with Lotka-Volterra dynamics. Physical Review E 95:042414. 10.1103/PhysRevE.95.042414

Case TJ (2000) An Illustrated Guide to Theoretical Ecology. Oxford University Press, New York

Cockrell C, O’Sullivan JD, Terry JCD, et al (2024) Self-organization of ecosystems to exclude half of all potential invaders. Physical Review Research 6:013093. 10.1103/PhysRevResearch.6.013093

Cottenie K (2005) Integrating environmental and spatial processes in ecological community dynamics. Ecology Letters 8(11):1175–1182. 10.1111/j.1461-0248.2005.00820.x

DeAngelis DL, Grimm V (2014) Individual-based models in ecology after four decades. F1000Prime Reports 6:39. 10.12703/P6-39

Driscoll DA, Lindenmayer DB (2010) Empirical tests of metacommunity theory using an isolation gradient. Ecological Monographs 79(3):485–501. 10.1890/08-1114.1

Grimm V, Railsback SF (2005) Individual-based Modeling and Ecology. Princeton Series in Theoretical and Computational Biology, Princeton University Press, Princeton, 10.1515/9781400850624

Hastings A, Botsford LW (2006) Persistence of spatial populations depends on returning home. Proceedings of the National Academy of Sciences 103(15):6067–6072. 10.1073/pnas.0506651103

Hofbauer J (1994) Heteroclinic cycles in ecological differential equations. In: Equadiff 8. Mathematical Institute, Slovak Academy of Sciences, pp 105–116, URL http://eudml.org/doc/220311

Hofbauer J, Sigmund K (1998) Evolutionary Games and Population Dynamics. Cambridge University Press, Cambridge

Holyoak M, Leibold MA, Holt RD (eds) (2005) Metacommunities: Spatial Dynamics and Ecological Communities. University of Chicago Press, Chicago

Jackson ZR, Leibold MA, Holt RD, et al (2025) Modeling and inferring metacommunity dynamics with Maximum Caliber. Proceedings of the National Academy of Sciences 122(28):e2520867123. 10.1073/pnas.2520867123

Kendall DG (1948) On the generalized ‘birth-and-death’ process. The Annals of Mathematical Statistics 19(1):1–15. URL http://www.jstor.org/stable/2236051

Kerr B, Riley MA, Feldman MW, et al (2002) Local dispersal promotes biodiversity in a real-life game of rock-paper-scissors. Nature 418(6894):171–174. 10.1038/nature00823

Kessler DA, Shnerb NM (2015) Generalized model of island biodiversity. Physical Review E 91(4):042705. 10.1103/PhysRevE.91.042705

Kurtz TG (1970) Solutions of ordinary differential equations as limits of pure jump Markov processes. Journal of Applied Probability 7(1):49–58. 10.2307/3212147

Leibold MA, Holyoak M, Mouquet N, et al (2004) The metacommunity concept: a framework for multi-scale community ecology. Ecology Letters 7(7):601–613. 10.1111/j.1461-0248.2004.00608.x

Leigh EGJ (1981) The average lifetime of a population in a varying environment. Journal of Theoretical Biology 90(2):213–239. 10.1016/0022-5193(81)90044-8

Lerch BA, Rudrapatna A, Rabi N, et al (2023) Connecting local and regional scales with stochastic metacommunity models: Competition, ecological drift, and dispersal. Ecological Monographs 93(4):e1591. 10.1002/ecm.1591

Logue JB, Mouquet N, Peter H, et al (2011) Empirical approaches to metacommunities: a review and comparison with theory. Trends in Ecology & Evolution 26(9):482–491. 10.1016/j.tree.2011.04.009

MacNeil MA, Graham NAJ, Polunin NVC, et al (2009) Hierarchical drivers of reeffish metacommunity structure. Ecology 90(1):252–264. 10.1890/07-0487.1

Mallmin E, Traulsen A, De Monte S (2024) Chaotic turnover of rare and abundant species in a strongly interacting model community. Proceedings of the National Academy of Sciences 121(11):e2312822121. 10.1073/pnas.2312822121

May RM, Leonard WJ (1975) Nonlinear aspects of competition between three species. SIAM Journal on Applied Mathematics 29(2):243–253. 10.1137/0129022

Moilanen A (2004) SPOMSIM: software for stochastic patch occupancy models of metapopulation dynamics. Ecological Modelling 179(4):533–550. 10.1016/j.ecolmodel.2004.04.019

Nwankwo EC, Rossberg AG (2026) Widespread slowdown in short-term species turnover despite accelerating climate change. Nature Communications 17:1450. 10.1038/s41467-025-68187-1

O’Sullivan JD, Knell RJ, Rossberg AG (2019) Metacommunity-scale biodiversity regulation and the self-organised emergence of macroecological patterns. Ecology Letters 22(9):1428–1438. 10.1111/ele.13294

O’Sullivan JD, Terry JCD, Rossberg AG (2021) Intrinsic ecological dynamics drive biodiversity turnover in model metacommunities. Nature Communications 12:3627. 10.1038/s41467-021-23769-7

O’Sullivan JD, Terry JCD, Rossberg AG (2023a) Temporally robust occupancy frequency distributions in riverine metacommunities explained by local biodiversity regulation. Global Ecology and Biogeography 32(12):2145–2158. 10.1111/geb.13756

O’Sullivan JD, Terry JCD, Wilson R, et al (2023b) Community composition exceeds area as a predictor of long-term conservation value. PLOS Computational Biology 19(1):e1010804. 10.1371/journal.pcbi.1010804

Ovaskainen O (2002) The effective size of a metapopulation living in a heterogeneous patch network. The American Naturalist 160(5):612–628. 10.1086/342818

Pärtel M, Tamme R, Carmona CP, et al (2025) Global impoverishment of natural vegetation revealed by dark diversity. Nature 641:917–924. 10.1038/s41586-025-08814-5

Arnoulx de Pirey T, Bunin G (2024) Many-species ecological fluctuations as a jump process from the brink of extinction. Physical Review X 14(1):011037. 10.1103/PhysRevX.14.011037

Reichenbach T, Mobilia M, Frey E (2007) Mobility promotes and jeopardizes biodiversity in rock-paper-scissors games. Nature 448(7157):1046–1049. 10.1038/nature06095

Ricklefs RE (2008) Disintegration of the ecological community. The American Naturalist 172(6):741–750. 10.1086/593002

Rossberg AG (2007) Some first principles of complex systems theory. RIMS Kôkyûroku 1551:129–136. URL http://axel.rossberg.net/paper/Rossberg2007d.pdf

Rossberg AG (2013) Food Webs and Biodiversity: Foundations, Models, Data. John Wiley & Sons, Chichester, 10.1002/9781118502181

Thompson PL, Rayfield B, Gonzalez A (2017) Loss of habitat and connectivity erodes species diversity, ecosystem functioning, and stability in metacommunity networks. Ecography 40(1):98–108. 10.1111/ecog.02558

Volterra V (1931) Leçons sur la théorie mathématique de la lutte pour la vie. Gauthier-Villars, Paris

Wang PC, Finley JC (2011) A landscape of shifting-mosaic steady state in Lassen Volcanic National Park, California. Ecological Research 26(1):191–199. 10.1007/s11284-010-0776-1

Winegardner AK, Jones BK, Ng ISY, et al (2012) The terminology of metacommunity ecology. Trends in Ecology & Evolution 27(5):253–254. 10.1016/j.tree.2012.01.007

