## Supplementary Information for "A modelling technique unifying four paradigms of metacommunity theory"

### Supplementary Information: Novel modelling technique unifies four paradigms of metacommunity theory

This supplement provides discussion of details on selected topics as referenced in the main text.

### 1 S1: ODE Model Implementation Details

#### 1.1 Numerical solution in log-space

For small immigration fluxes $I$, population biomasses $B_i$ can become extremely small, generating numerical difficulties with standard ODE solvers (e.g., spurious negative biomasses). To avoid this, we solve numerically the equivalent problem for the transformed variables $y_i = \ln B_i$, for which the general dynamics equation becomes

$$\frac{\mathrm{d}y_i}{\mathrm{d}t} = g_i(e^{y_1}, \ldots, e^{y_S}) + I_i\, e^{-y_i}. \tag{1}$$

This logarithmic transformation is standard for stiff systems with variables spanning many orders of magnitude (Press et al., 2007). We solve Eq. (1) using the variable-step, variable-order backward differentiation formula (BDF) integrator CVode from the SUNDIALS library (Hindmarsh et al., 2005) through the Python interface provided by the Assimulo package (Andersson et al., 2015).

### 2 S2: IBM Simulation Algorithm

#### 2.1 Update procedure

We simulate the IBM using a discrete-time approximation with fixed time step $\Delta t = 0.01$. At each time step, we update populations through three sequential operations. First, survival: each of the $N_i$ individuals survives with probability $\exp(-\mu_i \Delta t)$, where $\mu_i = \max(-g_i, M)$ is the effective mortality rate. The number of survivors is drawn from a binomial distribution

$$N_i^{\text{surv}} \sim \text{Binomial}(N_i, e^{-\mu_i \Delta t}). \tag{2}$$

Second, reproduction: surviving individuals give birth. The expected number of offspring per survivor over the interval is $e^{(g_i+\mu_i)\Delta t} - 1$, and the total births are drawn from a Poisson distribution

$$N_i^{\text{birth}} \sim \text{Poisson}\big((e^{(g_i+\mu_i)\Delta t} - 1) \cdot N_i^{\text{surv}}\big). \tag{3}$$

Third, immigration: new individuals arrive through the immigration flux

$$N_i^{\text{imm}} \sim \text{Poisson}\big(I_i \cdot \Delta t / m_0\big). \tag{4}$$

The updated population is $N_i \leftarrow N_i^{\text{surv}} + N_i^{\text{birth}} + N_i^{\text{imm}}$. This formulation ensures that the IBM dynamics matches both demographic variance of birth-death-immigration processes and, in expectation, the ODE dynamics to first order in $\Delta t$.

##### 2.2 Complete algorithm

**Algorithm 1** Individual-based model (IBM) simulation

**Require:** Intrinsic growth rates $r_i$, competition matrix $C_{ij}$, body mass $m_0$, immigration flux $I_i$, baseline mortality $M$, time step $\Delta t$, total steps $n_{\text{steps}}$

Initialise $N_i \leftarrow \lfloor 0.1/m_0 \rfloor$ for all species $i$ ▷ Initial population
**for** $s = 1$ to $n_{\text{steps}}$ **do**
  $B_i \leftarrow N_i \cdot m_0$ for all $i$ ▷ Convert to biomass
  $g_i \leftarrow r_i - \sum_j C_{ij} B_j$ for all $i$ ▷ Net growth rate
  $\mu_i \leftarrow \max(M, -g_i)$ for all $i$ ▷ Effective mortality
  $N_i \leftarrow \text{Binomial}(N_i, e^{-\mu_i \Delta t})$ ▷ Survival
  $\lambda_i^{\text{birth}} \leftarrow (e^{(g_i+\mu_i)\Delta t} - 1) \cdot N_i$
  $N_i \leftarrow N_i + \text{Poisson}(\lambda_i^{\text{birth}})$ ▷ Births
  $N_i \leftarrow N_i + \text{Poisson}(I_i \Delta t / m_0)$ ▷ Immigration
**end for**
**return** Trajectory $\{N_i(t) \cdot m_0\}$

### 3 S3: PSD Derivation and Implementation

We detail the derivation of the Probabilistic–Stochastic–Deterministic approximation scheme.

#### 3.1 Derivation of S-state dynamics

##### 3.1.1 Poisson clocks

We begin by deriving the dynamics of the Poisson clocks. First, observe that the natural logarithm of a uniformly distributed random variate $U$ is $-1$ times an exponentially distributed random variate $X$ with mean 1. To verify this, observe for any $0 < u < 1$ we have $\Pr[U < u] = u$ and, with $x$ defined by $x = -\log(u)$,

$$\Pr[X > x] = \Pr[-X < -x] = \Pr[\log(U) < \log(u)] = \Pr[U < u] = u = \exp(-x). \tag{5}$$

Hence, initial values of Poisson clocks $P_i(t)$ time $-1$ follow a unit exponential distribution.

The dynamics of Poisson clocks in the PSD algorithm is such that their values can never decrease. With this provision, the probability distribution of a Poisson clock conditional to not having passed zero yet is time invariant. To see this, let $P_i(t)$ be a Poisson clock, $X$ be the exponentially distributed initial value of $-P_i(t)$ and $\Delta P \geq 0$ the amount by which the Poisson clock has dynamically increased so far. We evaluate the distribution of $-P_i(t) = X - \Delta P$ conditional to $P_i(t) = -X + \Delta P$ remaining smaller than zero, in order to verify that this remains an exponential distribution. With $x > 0$, some algebraic re-arrangement and Eq. (5), we do indeed obtain the conditional probability $\Pr[X - \Delta P > x | -X + \Delta P < 0]$ as

$$\frac{\Pr[X - \Delta P > x]}{\Pr[-X + \Delta P < 0]} = \frac{\Pr[X > x + \Delta P]}{\Pr[-X < -\Delta P]} = \frac{\Pr[X > x + \Delta P]}{\Pr[X > \Delta P]} = \frac{\exp(-x - \Delta P)}{\exp(-\Delta P)} = \exp(-x). \tag{6}$$

As a result, the probability that a Poisson clock $P_i(t)$ that has not yet run out reaches zero within a short time interval $t$ to $t + \Delta t$ is, to lowest order in $\Delta t$, given by

$$\Pr\left[P_i(t) + \frac{\mathrm{d}P_i(t)}{\mathrm{d}t}\Delta t \geq 0\right] = \Pr\left[P_i(t) \geq -\frac{\mathrm{d}P_i(t)}{\mathrm{d}t}\Delta t\right] =$$
$$1 - \Pr\left[-P_i(t) > \frac{\mathrm{d}P_i(t)}{\mathrm{d}t}\Delta t\right] = 1 - \exp\left(-\frac{\mathrm{d}P_i(t)}{\mathrm{d}t}\Delta t\right) = \frac{\mathrm{d}P_i(t)}{\mathrm{d}t}\Delta t + \text{h.o.t}, \quad (7)$$

where h.o.t denotes highter-order terms in $\Delta t$.

For ecological consistency, we therefore need to advance Poisson clocks of species in S-states at a rate that equals the rate at which individuals arrive and establish new populations. The arrival rate is $I_i/m_i$. Demographic stochasticity can cause extirpation of a population in the early phase of population growth, preventing its establishment even when the population growth rate $g_i$ is positive. To obtain the establishment probability, we make the simplifying assumptions that birth rate $\beta_i = g_i + \mu_i$ and death rates $\mu_i$ are both constant over the time interval over which establishment is determined. In particular, we assume that at by time population sizes are so large that density dependence becomes relevant stochastic extinction has become virtually impossible. We can then rely on the well-known result for simple density-independent birth-death processes (Kendall, 1948) that

$$p_i^* = \Pr[\text{establish}] = 1 - \frac{\mu_i}{\beta_i} = \frac{\beta_i - \mu_i}{\beta_i} = \frac{g_i}{g_i + \mu_i}. \quad (8)$$

Putting it all together, noting that $\mu_i = M_i$ for $g_i$ close to zero, and using the approximation $g_i \approx \hat{g}_i$ to assure internal consistency of the PSD approach leads to the formula

$$\frac{\mathrm{d}P_i}{\mathrm{d}t} = \frac{I_i}{m_i} \cdot \frac{\hat{g}_i}{\hat{g}_i + M_i}. \quad (9)$$

##### 3.1.2 Biomass of failed colonisers

To derive an approximation for the dynamics of expected population biomass in S-states conditional to a failure of that population to establish, our starting point is the unconditional probability generating function $G(s,t)$ for a randomly varying population size $N(t)$, defined as the expectation value of $s^{N(t)}$, that is,

$$G(s,t) = \sum_{n=0}^{\infty} \Pr[N(t) = n]\, s^n, \quad (10)$$

where $\Pr[N(t) = n]$ is the probability that the population size at time $t$ is $n$. It is known that for a population that had size 1 at $t = 0$ and since then evolved according to a simple birth-death process with fixed rates of birth $\beta$ and death $\mu$ and $\beta > \mu$,

$$G(s,t) = \frac{(s-1)(\beta - \mu)}{(\beta s - \mu)e^{t(\mu - \beta)} - \beta s + \beta} + 1. \quad (11)$$

Noting the structure of $G(s,t)$ given in Eq. (10), the probability of extirpation of that population can be obtained from this as

$$\Pr[\text{extinct}] = \lim_{t \to \infty} G(0,t) = \frac{\mu}{\beta}, \quad (12)$$

consistent with Eq. (8). Furthermore, we recall for later reference that, by standard arguments drawing on Eq. (10), the expected population size at time $t$ can be computed from $G(s,t)$ as

$$\left.\frac{\partial G(s,t)}{\partial s}\right|_{s=1} = e^{t(\beta-\mu)} = e^{gt}, \tag{13}$$

where we introduced the growth rate $g = \beta - \mu$ in the last step.

Now we restrict this process to those cases where population eventually does go extinct. For any time $t > 0$, the probability that the population has $n$ individuals at that time and later goes extinct (requiring that the lineages founded by each of these $n$ individuals eventually go extinct) is

$$\Pr[N(t) = n] \cdot (\Pr[\text{extinct}])^n = \Pr[N(t) = n] \left(\frac{\mu}{\beta}\right)^n. \tag{14}$$

Using this and Eq. (10), we can therefore obtain the probability generating function of the population size at time $t$ conditional to the eventual extirpation of that population as

$$G_{\text{ex}}(s,t) = \frac{G(s\,\mu/\beta, t)}{G(\mu/\beta, t)} = \frac{\beta(s-1)e^{t(\mu-\beta)} + \beta - \mu s}{\mu(s-1)e^{t(\mu-\beta)} + \beta - \mu s}. \tag{15}$$

By the same approach as in Eq. (13), we then obtain the expected population size at time $t$ conditional to the eventual extirpation of the population as

$$\left.\frac{\partial G_{\text{ex}}(s,t)}{\partial s}\right|_{s=1} = e^{t(\mu-\beta)} = e^{-gt}, \tag{16}$$

with the sign of $g$ reversed compared to Eq. (13). Approximating again $g_i \approx \hat{g}_i$ for internal consistency of the PSD formalism and assuming that $\beta$ and $\mu$ won't change much while populations attempt but fail to establish, we therefore approximate the dynamics of expected population size while establishment in the S-state is attempted but fails as

$$\frac{\mathrm{d}B_i}{\mathrm{d}t} = -\hat{g}_i\, B_i + I_i. \tag{17}$$

By tracking this expected biomass we can capture the effect of cases where populations manage to disperse before they fail to establish. We also obtain valid estimates of expected biomass at instances where S-state populations transition to P-states.

##### 3.2 Transition probability derivation

When $\hat{g}_i$ crosses zero from below, propagules already present may or may not establish. The approximation of constant birth and death rates used above would predict that at the zero crossing, with $\beta = \mu$, no propagule establishes by Eq. (12). In fact, however, zero crossings resulting from the re-organisation of communities after immigration of other species can be rather fast, which is why a more refined approach is needed.

We therefore estimate the probability of establishment using the theory of birth-death processes for time-dependent parameters by Branson (1991). We assume that the rate $c \geq 0$ at which $g_i$ changes during the time period determining establishment is fixed and choose the time axis such that the zero crossing occurs exactly when $t = 0$. For our model, this implies

$$\beta = M + c\,t \tag{18a}$$

and

$$\mu = M. \tag{18b}$$

The theory of Branson (1991) requires the auxiliary function defined as

$$\rho(T,t) = \int_T^t [\mu(\tau) - \beta(\tau)]d\tau, \tag{19}$$

which, by Eq. (18), evaluates here to $\rho(T,t) = c(T^2 - t^2)/2$. Evaluating further, in this order, Branson's Eqs. (14) and (13), and then Eq. (6), we obtain the probability $P_0(T,t)$ of extirpation until $t$ of a population seeded with a single individual at $T$ as

$$P_0(T,t) = 1 - \left\{ \sqrt{\frac{\pi}{2c}} M \exp\left(\frac{cT^2}{2}\right) \left[ \operatorname{erf}\left(\frac{c^{1/2}t}{\sqrt{2}}\right) - \operatorname{erf}\left(\frac{c^{1/2}T}{\sqrt{2}}\right)\right] + 1 \right\}^{-1}. \tag{20}$$

We choose the starting point at $T = 0$ because at this point the population potentially transitions into the S-state, where the PSD formalism accounts entirely differently for immigations than in the P-state. The probability of extirpation asympototes for large $t$ to

$$P_x = \lim_{t\to\infty} P_0(0,t) = 1 - \left(\sqrt{\frac{\pi}{2c}} M + 1\right)^{-1} = 1 - \frac{2c^{1/2}}{2c^{1/2} + \sqrt{2\pi} M}. \tag{21}$$

To apply this result within the PSD approximation, recall that in the P-state, corresponding to time $t \leq 0$ with the time axis chosen as above, the actual number of individuals present is not known, only the expectation value $B_i/m_i$ of that number. In order to obtain from Eq. (21) a model for this situation, we assume population size to be Poisson distributed for simplicity. The known probability generating function of a Poisson distribution with mean $\lambda \geq 0$ is $G_{\text{Pois}}(s,\lambda) = \exp(\lambda(s-1))$. By formal arguments analogous to those used in Sec. 3.1, we obtain from this and Eq. (21) immediately the approximate probability that the population does not establish as

$$P(\text{P} \to \text{S}) = G_{\text{Pois}}\left(P_x, \frac{B_i}{m_0}\right) = \exp\left(-\frac{B_i}{m_0} \cdot \frac{2c^{1/2}}{2c^{1/2} + \sqrt{2\pi}\, M_i}\right), \tag{22}$$

where $B_i$ is the expected population biomass at the zero crossing of $\hat{g}_i$ and $c > 0$ is the rate of change of $\hat{g}_i$ at the time of the zero crossing. In the limit $c \to 0$, Eq. (22) predicts eventual extirpation of all population present before the tansition point. In the limit $c \to \infty$, it predicts that any population present before the transition point will establish. In the latter case, the question whether establishment and so a direct transtition from P- to D-state occurs depends only on whether a population was present at the time of the transition at all.

Future research might find more accurate analytic approximations of the probability generating function of population size at the transition point, permitting refinement of Eq. (22).

#### 3.3 Biomass adjustments at transitions

In simple birth-death processes, expectation values of population sizes strictly follow the corresponding ODE equation. To maintain consistency with this expected-value dynamics, we adjust population biomasses in state stransitions as follows:

P → D transition: The biomass $B_i$ is divided by the probability that this transition occurs, i.e., by $1 - P(\mathrm{P} \to \mathrm{S})$:

$$B_i \leftarrow \min\left(\frac{B_i}{1 - P(\mathrm{P} \to \mathrm{S})}, B_{\mathrm{cap}}\right), \tag{23}$$

where $B_{\mathrm{cap}} = 1$ is a heuristic cap preventing unrealistic values when $P(\mathrm{P} \to \mathrm{S})$ is close to 1.

S → D transition: The biomass is that the one individual that arrived as the Poisson clock ran out, divided by the establishment probability:

$$B_i \leftarrow \min\left(\frac{M_i}{\hat{g}_i/(\hat{g}_i + \mu_i)}, B_{\mathrm{cap}}\right) = \min\left(\frac{M_i(\hat{g}_i + \mu_i)}{\hat{g}_i}, B_{\mathrm{cap}}\right). \tag{24}$$

In both cases, the adjustments, while introducing an un-ecological discontinuity in at the moment of the state transition, become valid after sufficient time has passed such that lineages that failed to establish a resident population have locally died out.

##### 3.4 Numerical implementation

The PSD framework can be implemented using two different numerical strategies, each suited to different simulation scenarios.

For single-patch simulations and small metacommunities where state transitions (P → S, S → D, D → P) are relatively infrequent, we employ an event-driven approach. This implementation uses the SUNDIALS CVode library (Hindmarsh et al., 2005), which provides a variable-step, variable-order backward differentiation formula (BDF) integrator with adaptive step-size control. The adaptive algorithm automatically adjusts the integration step to maintain accuracy while taking large steps during periods of slow dynamics. Crucially, CVode includes root-finding capabilities that detect precisely when continuous variables cross specified thresholds. We use this functionality to identify the exact moments when the invasion growth rate $\hat{g}_i$ crosses zero (triggering P ↔ S or S ↔ D transitions) or when a Poisson clock $P_i$ reaches zero (triggering S → D establishment). This event-driven approach is computationally efficient when transitions are rare, as the solver can take large time steps between events.

For large, species-rich metacommunities where state transitions occur frequently across many patches and species, we instead use a fixed-step forward Euler scheme. At each time step $\Delta t$, we update all continuous variables (biomasses $B_i$ and Poisson clocks $P_i$) and then test for sign changes in $\hat{g}_i$ and threshold crossings in $P_i$ to detect transitions. Although this approach lacks the precision of event-driven root-finding and requires smaller time steps for accuracy, it avoids the computational overhead of stopping and re-starting ODE simulations at each separate event. The simulations presented in this paper use $\Delta t = 0.01$, which provides sufficient accuracy for the parameter regimes explored. We would welcome suggestions for improved implementations that efficiently handle frequent events occuring over large metacommunities while retaining the accuracy and speed of modern variable-step-size ODE solvers.

##### 3.5 Complete algorithm

Algorithm 2 presents the fixed-step Euler implementation in pseudocode form. The event-driven implementation follows the same logical structure but replaces the discrete time-stepping loop with quasi-continuous integration punctuated by root-finding events.

**Algorithm 2** Probabilistic–Stochastic–Deterministic (PSD) simulation (fixed-step)

**Require:** Intrinsic growth rates $r_i$, competition matrix $C_{ij}$, body mass $m_0$, immigration flux $I$, baseline mortality $M$, time step $\Delta t$, total steps $n_{\text{steps}}$, biomass cap $B_{\text{cap}} = 1$

**Initialise:**
  $B_i \leftarrow m_0/10$ for all species $i$
  $\text{state}_i \leftarrow \text{S}$ for all $i$ ▷ All species start in waiting state
  $P_i \leftarrow \ln(U_i)$ where $U_i \sim \text{Uniform}(0, 1)$ ▷ Initialise Poisson clocks
**for** $s = 1$ to $n_{\text{steps}}$ **do**
  ▷ **Step 1: Compute growth rates**
  $g_i \leftarrow r_i - \sum_j C_{ij} B_j$ for all $i$ ▷ All species contribute to competition
  $\hat{g}_i \leftarrow g_i + C_{ii} B_i$ for all $i$

  ▷ **Step 2: Handle state transitions**
  **for** each species $i$ **do**
    **if** $\text{state}_i = \text{D}$ **and** $\hat{g}_i < 0$ **then** ▷ D $\rightarrow$ P
      $\text{state}_i \leftarrow \text{P}$
    **else if** $\text{state}_i = \text{P}$ **and** $\hat{g}_i \geq 0$ **then** ▷ P $\rightarrow$ S or P $\rightarrow$ D
      $c_i \leftarrow |\Delta \hat{g}_i / \Delta t|$
      $p_{\text{S}} \leftarrow \exp\left(-\frac{B_i}{m_0} \cdot \frac{2\sqrt{c_i}}{2\sqrt{c_i} + \sqrt{2\pi}\, M}\right)$
      Draw $u \sim \text{Uniform}(0, 1)$
      **if** $u < p_{\text{S}}$ **then**
        $\text{state}_i \leftarrow \text{S}$; $P_i \leftarrow \ln(U)$ where $U \sim \text{Uniform}(0, 1)$
      **else**
        $\text{state}_i \leftarrow \text{D}$; $B_i \leftarrow \min(B_i/(1 - p_{\text{S}}), B_{\text{cap}})$
      **end if**
    **else if** $\text{state}_i = \text{S}$ **and** $\hat{g}_i < 0$ **then** ▷ S $\rightarrow$ P
      $\text{state}_i \leftarrow \text{P}$
    **end if**
  **end for**

  ▷ **Step 3: Update dynamics**
  **for** each species $i$ **do**
    **if** $\text{state}_i = \text{D}$ **or** $\text{state}_i = \text{P}$ **then**
      $\ln B_i \leftarrow \ln B_i + (g_i + I/B_i) \cdot \Delta t$
      $B_i \leftarrow \min(B_i, B_{\text{cap}})$
    **else if** $\text{state}_i = \text{S}$ **then**
      $p_i^* \leftarrow \hat{g}_i / (\hat{g}_i + M)$
      $B_i \leftarrow B_i + (-\hat{g}_i \cdot B_i + I) \cdot \Delta t$
      $P_i \leftarrow P_i + (I/m_0) \cdot p_i^* \cdot \Delta t$
      **if** $P_i \geq 0$ **then** ▷ S $\rightarrow$ D: Establishment
        $\text{state}_i \leftarrow \text{D}$
        $B_i \leftarrow \min(M/p_i^*, B_{\text{cap}})$
        $P_i \leftarrow \ln(U)$ where $U \sim \text{Uniform}(0, 1)$
      **end if**
    **end if**
  **end for**
**end for**
**return** Trajectory $\{B_i(t)\}$ and state history

### 4 S4: Parameter Values

Table 1 summarises the parameter values used in single-patch simulations.

Table 1: **Default parameter values for single-patch simulations.**

| **Parameter** | **Symbol** | **Value** | **Description** |
|---|---|---|---|
| Species number | $S$ | 300 | Number of species |
| Intrinsic growth rate | $r_i$ | 1 | All species |
| Intraspecific competition | $C_{ii}$ | 1 | All species |
| Interaction strength | | 0.4 | Non-zero off-diagonal $C_{ij}$ |
| Connectance | | 0.4 | Probability of interaction |
| Baseline mortality | $M$ | 0.2 | All species |
| Immigration flux | $I$ | $10^{-8}$ | Biomass per unit time |
| Body mass (low) | $m_0$ | $10^{-11}$ | Many small individuals |
| Body mass (intermediate) | $m_0$ | $10^{-7.6}$ | Transition regime |
| Body mass (high) | $m_0$ | $10^{-4}$ | Few large individuals |
| Time step | $\Delta t$ | 0.01 | Simulation step |
| Simulation time | $T$ | $10^4$ | Total duration |
| Biomass cap | $B_{\text{cap}}$ | 1 | Maximum biomass |

### 5 S5: Single-patch Validation — Additional Details

#### 5.1 Simulation protocol

We simulated communities of $S = 500$ species using identical parameters across all three models: intrinsic growth rates $r_i = 1$, with interaction matrix sampled as described in Section S1. A constant immigration flux $I = 10^{-8}$ was applied uniformly to all species. For the ODE and PSD models, we used an event-handling ODE solver with variable step-size integration via the SUNDIALS CVode library (see Section S3), while the IBM used a fixed time step of $\Delta t = 0.01$.

To explore the role of demographic stochasticity, we compared three body-mass regimes. At low body mass ($m_0 = 10^{-11}$), populations consist of numerous small individuals where stochastic effects are minimal due to the law of large numbers. At high body mass ($m_0 = 10^{-4}$), populations contain few large individuals where demographic noise is pronounced and can qualitatively alter community dynamics. We also tested an intermediate body mass ($m_0 = 10^{-7.6}$), corresponding to the transition point where $m_0 \approx B_{\text{lower}}$, to verify that PSD accurately captures IBM dynamics across the full parameter range.

Each simulation ran for $T = 10^4$ time units, corresponding to $10^6$ discrete update steps for the IBM.

#### 5.2 Statistical analysis

Summary statistics were computed after discarding the initial 20% of each simulation as burn-in to remove transient effects. Species richness was defined as the number of species with biomass $B_i > 10^{-3}$ at any given time point; we report the mean richness over the post-burn-in period.

Invasion rate quantifies how frequently new species successfully colonise the community. We define a successful invasion as an event where a species crosses upward through a biomass threshold of $B_{\text{th}} = 10^{-3}$: that is, a species with $B_i(t) < B_{\text{th}}$ at time $t$ transitions to $B_i(t + \Delta t) > B_{\text{th}}$ at the next recorded time point. The invasion rate is then the total number of such threshold-crossing events divided by the elapsed time. To estimate uncertainty, we divided the post-burn-in time series into $n = 10$ non-overlapping chunks, computed the invasion rate within each chunk, and report the mean and standard error across chunks.

Model comparisons used one-way ANOVA to test for overall differences among the three models (ODE, IBM, PSD). When ANOVA indicated significant differences ($p < 0.05$), we applied Tukey's Honestly Significant Difference (HSD) test for pairwise comparisons to identify which specific model pairs differed.

### 6 S6: Multi-patch Metacommunity — Model Details

#### 6.1 Rock–paper–scissors competition

For the rock–paper–scissors (RPS) system, we consider $S = 3$ species with cyclic competitive interactions. The competition matrix takes the form

$$\mathbf{C} = \begin{pmatrix} 1 & \alpha & \beta \\ \beta & 1 & \alpha \\ \alpha & \beta & 1 \end{pmatrix}, \qquad \alpha > 1 > \beta \geq 0, \tag{25}$$

where intraspecific competition is normalised to unity. We set intrinsic growth rates $r_i = 1$ for all species. The dominant off-diagonal parameter $\alpha$ then establishes the cyclic competitive hierarchy: species 1 suppresses species 3, species 2 suppresses species 1, and species 3 suppresses species 2. In numerical examples, we use interaction strengths $\alpha = 1.7$ and $\beta = 0.4$, which places the system in the oscillatory regime where cyclic dominance produces sustained oscillations in species abundances.

#### 6.2 Spatial grid and boundary conditions

Patches are arranged on an $L_x \times L_y$ grid with periodic (toroidal) boundary conditions, eliminating edge effects. For the simulations presented in the main text, we use a $20 \times 20$ grid (400 patches total). The model state becomes $B_{i,\mathbf{x}}(t)$ for species $i$ at patch $\mathbf{x} = (x, y)$.

#### 6.3 Dispersal kernels

During each time step, a fraction of biomass leaves each patch and is redistributed to other patches according to a dispersal kernel. We consider two limiting dispersal modes:

##### 6.3.1 Local dispersal

Under local dispersal, propagules move only to immediately adjacent patches. The flux from patch $\mathbf{x}$ to patch $\mathbf{y}$ is

$$\Phi_i(\mathbf{x} \to \mathbf{y}) = \begin{cases} \dfrac{D}{4}\, B_{i,\mathbf{x}}, & \mathbf{y} \in \mathcal{N}(\mathbf{x}), \\ 0, & \text{otherwise}, \end{cases} \tag{26}$$

where $\mathcal{N}(\mathbf{x})$ denotes the four von Neumann neighbours of patch $\mathbf{x}$ (north, south, east, west) and $D$ is the dispersal rate (fraction of biomass leaving per unit time). This represents organisms with limited mobility, such as plants with gravity-dispersed seeds or invertebrates with crawling larvae.

###### 6.3.2 Global dispersal

Under global (long-distance) dispersal, propagules can reach any patch in the metacommunity with equal probability. The flux becomes

$$\Phi_i(\mathbf{x} \to \mathbf{y}) = \frac{D}{L_x L_y} B_{i,\mathbf{x}}, \qquad \forall\, \mathbf{y} \neq \mathbf{x}, \tag{27}$$

representing organisms with long-range dispersal capabilities, such as wind-dispersed spores, migratory birds, or species subject to anthropogenic transport.

###### 6.3.3 Immigration flux

Under either dispersal mode, the total immigration flux into patch $\mathbf{y}$ is the sum of contributions from all source patches:

$$I_i(\mathbf{y}, t) = \sum_{\mathbf{x}\neq\mathbf{y}} \Phi_i(\mathbf{x} \to \mathbf{y}) = \sum_{\mathbf{x}\neq\mathbf{y}} K_{\mathbf{xy}}\, B_{i,\mathbf{x}}(t), \tag{28}$$

where $K_{\mathbf{xy}}$ is the dispersal kernel connecting patches $\mathbf{x}$ and $\mathbf{y}$. We do not explicitly account for local mass loss due to dispersal, as in our model it would simply amount to a uniform correction to in intrinsic growth rates $r_i$.

For the spatial pattern formation simulations presented in Section 5 of the main text, we use global dispersal (Eq. 27) to maximise spatial coupling and reveal the full range of collective dynamics.

##### 6.4 Simulation protocol

We simulated the RPS metacommunity on a $20\times20$ grid using all three modelling frameworks (ODE, IBM, and PSD) with identical parameters and initial conditions. Each patch was initially assigned to one of the three species with equal probability ($p = 1/3$ each), introducing the spatial heterogeneity that seeds pattern development. Simulations ran for $T$ = 100,000 unit times, sufficient for the system to develop and display characteristic spatio-temporal dynamics. The spatial distribution of dominant species was recorded at regular intervals ($\Delta t_{\text{output}}$ = 10,000) to visualise the evolution of spatial patterns.

##### 6.5 Parameter values for RPS simulations

#### 7 S7: Derivation of Metacommunity Oscillation Period

##### 7.1 Mean-field dynamics under global dispersal

Under pure long-distance dispersal, all patches receive equal immigration flux from the entire metacommunity. Let $P_i(t)$ denote the fraction of patches where species $i$ is the dominant (established) species. The constraint $P_1 + P_2 + P_3 = 1$ confines dynamics to a simplex.

The PSD waiting-state mechanism implies that patch fractions evolve according to invasion rates. In the rock–paper–scissors system, each species can only invade patches dominated by its inferior competitor: species 1 invades patches dominated by species 3, species 2 invades patches dominated by species 1, and species 3 invades patches dominated by species 2.

Table 2: **Parameter values for RPS metacommunity simulations.**

| **Parameter** | **Symbol** | **Value** | **Description** |
|---|---|---|---|
| Number of species | $S$ | 3 | Rock–paper–scissors |
| Grid size | $N_x \times N_y$ | $20 \times 20$ | 400 patches |
| Intrinsic growth rate | $r_i$ | 1 | All species |
| Dominant interaction | $\alpha$ | 1.7 | Cyclic hierarchy strength |
| Subdominant interaction | $\beta$ | 0.4 | Reverse interaction strength |
| Dispersal rate | $D$ | $5 \times 10^{-7}$ | Global dispersal |
| Body mass | $m_0$ | $10^{-4}$ | Per individual |
| Baseline mortality | $M$ | 0.2 | All species |
| Simulation time | $T$ | $10^5$ | Total duration |
| Time step | $\Delta t$ | 0.01 | Integration step |

##### 7.2 Transition rates

Consider the transition rate from a patch dominated by species $j$ to dominance by species $i$, where $i$ beats $j$ in the RPS hierarchy. Under the PSD framework, this rate is

$$\lambda_{j \to i} = \frac{D}{m_0} \cdot p_i^* \cdot P_i, \tag{29}$$

where $D$ is the dispersal rate (biomass emigrating per unit time per individual), $m_0$ is the body mass per individual, $p_i^*$ is the establishment probability of species $i$, and $P_i$ is the fraction of source patches containing species $i$. The factor $D/m_0$ represents the arrival rate of propagules (number of individuals per unit time), and $p_i^*$ is the probability that an arriving propagule successfully establishes.

##### 7.3 Establishment probability

For the symmetric RPS system with intrinsic growth rate $r_i = r$ and competition matrix from Eq. (25), we compute the establishment probability when species $i$ invades a patch dominated by species $j$ (where $i$ beats $j$).

In a patch dominated by species $j$, the resident population is at the single-species equilibrium $B_j^* = r/C_{jj}$. Because species $i$ beats species $j$, the competition coefficient from $j$ to $i$ is the subdominant value $\beta$ (the weak interaction in the direction opposite to the competitive hierarchy). The invader's growth rate is therefore

$$\hat{g}_i = r - C_{ij} B_j^* = r - \beta r = r(1 - \beta). \tag{30}$$

Using the establishment probability from the PSD framework (Eq. 6 in main text):

$$p^* = \frac{\hat{g}_i}{\hat{g}_i + M} = \frac{r(1-\beta)}{r(1-\beta) + M}, \tag{31}$$

where $M$ is the baseline mortality rate. For our parameter values ($r = 1$, $\beta = 0.4$, $M = 0.2$), this gives $p^* = 0.6/(0.6 + 0.2) = 0.75$.

#### 7.4 Mean-field dynamic equations

The dynamics of patch fractions follow:

$$\frac{dP_1}{dt} = \lambda_{3\to1}P_3 - \lambda_{1\to2}P_1 = \frac{D \cdot p^*}{m_0}(P_1P_3 - P_2P_1), \tag{32}$$

$$\frac{dP_2}{dt} = \lambda_{1\to2}P_1 - \lambda_{2\to3}P_2 = \frac{D \cdot p^*}{m_0}(P_2P_1 - P_3P_2), \tag{33}$$

$$\frac{dP_3}{dt} = \lambda_{2\to3}P_2 - \lambda_{3\to1}P_3 = \frac{D \cdot p^*}{m_0}(P_3P_2 - P_1P_3). \tag{34}$$

These equations have a symmetric equilibrium at $P_1^* = P_2^* = P_3^* = 1/3$.

#### 7.5 Linearisation and eigenvalue analysis

Using the constraint $P_3 = 1 - P_1 - P_2$, we reduce to two independent variables. Linearising around the symmetric equilibrium $(P_1, P_2) = (1/3, 1/3)$, the Jacobian matrix is:

$$\mathbf{J} = \frac{D \cdot p^*}{3m_0}\begin{pmatrix} -1 & -2 \\ 2 & 1 \end{pmatrix}. \tag{35}$$

The matrix $\mathbf{J}$ has purely imaginary eigenvalues:

$$\lambda_\pm = \frac{D \cdot p^*}{3m_0}\left(\pm i\sqrt{3}\right), \tag{36}$$

giving the angular oscillation frequency:

$$\omega = \frac{\sqrt{3} \cdot D \cdot p^*}{3m_0} = \frac{D \cdot p^*}{\sqrt{3} \cdot m_0}. \tag{37}$$

#### 7.6 Period formula

The oscillation period is

$$T_{\text{meta}} = \frac{2\pi}{\omega} = \frac{2\sqrt{3}\,\pi \cdot m_0}{D \cdot p^*}. \tag{38}$$

Substituting the establishment probability $p^* = r(1-\beta)/(r(1-\beta)+M)$ from Eq. (31)

$$T_{\text{meta}} = \frac{2\sqrt{3}\,\pi \cdot m_0 \cdot (r(1-\beta)+M)}{D \cdot r(1-\beta)}, \tag{39}$$

which is Eq. 7 in the main text.

#### 7.7 Isolated-patch dynamics (ODE prediction)

For comparison, the isolated-patch RPS system follows Lotka–Volterra dynamics

$$\frac{\mathrm{d}B_i}{\mathrm{d}t} = B_i\left(r_i - \sum_j C_{ij}B_j\right), \tag{40}$$

where $r_i = 1$ for all species. With the competition matrix $\mathbf{C}$ from Eq. (25), the coexistence equilibrium is $B_i^* = 1/(1+\alpha+\beta)$ for all species.

Linearising around this equilibrium, the Jacobian has eigenvalues with imaginary part (May and Leonard, 1975; Hofbauer and Sigmund, 1998)

$$\omega_{\text{isolated}} = \frac{\sqrt{3}\,(\alpha - \beta)}{2(1 + \alpha + \beta)}, \tag{41}$$

giving period

$$T_{\text{isolated}} = \frac{4\pi(1 + \alpha + \beta)}{\sqrt{3}\,(\alpha - \beta)}. \tag{42}$$

Note that $T_{\text{isolated}}$ depends only on competition parameters $(\alpha, \beta)$ and is independent of body mass $m_0$ and dispersal rate $D$.

##### 7.8 Numerical values

For the parameters used in the main text ($r = 1$, $\alpha = 1.7$, $\beta = 0.4$, $M = 0.2$, $m_0 = 10^{-4}$, $D = 5 \times 10^{-7}$), substituting into the formulae derived above yields an establishment probability of $p^* = 0.75$, a metacommunity oscillation period of $T_{\text{meta}} \approx 2900$ time units, and an isolated-patch oscillation period of $T_{\text{isolated}} \approx 17$ time units. The metacommunity period is therefore approximately 170 times longer than the isolated-patch prediction, consistent with the numerical simulations presented in Figure 4 of the main text.

### References

Andersson C, Führer C, Åkesson J (2015) Assimulo: A unified framework for ODE solvers. Mathematics and Computers in Simulation 116:26–43. https://doi.org/10.1016/j.matcom.2015.04.007

Branson D (1991) Inhomogeneous birth-death and birth-death-immigration processes and the logarithmic series distribution. Stochastic Processes and their Applications 39(1):131–137. https://doi.org/10.1016/0304-4149(91)90037-D

Hindmarsh AC, Brown PN, Grant KE, et al (2005) SUNDIALS: Suite of nonlinear and differential/algebraic equation solvers. ACM Transactions on Mathematical Software 31(3):363–396. https://doi.org/10.1145/1089014.1089020

Hofbauer J, Sigmund K (1998) Evolutionary Games and Population Dynamics. Cambridge University Press, Cambridge

Kendall DG (1948) On the generalized 'birth-and-death' process. The Annals of Mathematical Statistics 19(1):1–15. URL http://www.jstor.org/stable/2236051

May RM, Leonard WJ (1975) Nonlinear aspects of competition between three species. SIAM Journal on Applied Mathematics 29(2):243–253. https://doi.org/10.1137/0129022

Press WH, Teukolsky SA, Vetterling WT, et al (2007) Numerical Recipes: The Art of Scientific Computing, 3rd edn. Cambridge University Press, Cambridge
